# Transplantation of a kidney with a ureter and part of the bladder as a single block: an experimental study

**DOI:** 10.1101/2024.05.28.596363

**Authors:** Gani Kuttymuratov, Ardak Ainakulov, Askar Ayaganov, Kuat Oshakbayev, Arman Mirmanov, Daulet Zharasov, Zhandos Imanberdiev, Askar Taszhurekov, Bakhytzhan Abdimazhitov, Aruzhan Asanova, Tleuzhan Abdurakhman, Nurlybek Uderbayev, Arnagul Kalieva

## Abstract

**Objective:** To evaluate the effectiveness of en bloc transplantation of a donor kidney, ureters and part of the bladder to a recipient with simulated microcystis in an experimental trial.

**Methods:** Study Design: a 29-day, open, pilot prospective experimental trial: 14 days constituted an adaptation period, 5 days for the interventions, and 10 days constituted an observation period. The study totally included ten White Landras sibling pigs, which were divided into 12 donors and 12 recipients. The pigs were 3-4 months old and weighing 35-45 kg of both sexes from the same sow to avoid transplant rejection. The pigs lived 7-9 days after transplantation before they were euthanized, and then there were performed macrovisual and histological investigations. Descriptive, inferential statistics, and calculation of percentages were used. The Local Ethics Committee of West Kazakhstan Medical University approved the study.

**Results:** Eleven pigs survived the operation, but one pig died 10 hours after the operation. The cause of death was pulmonary embolism according to the pathological autopsy. In the eleven animals the kidney, ureters and part of the bladder transplanted as en block visually were filled with urine, full of blood, and tissue turgor was good. Visual inspection of the kidney and ureters was satisfactory, bright red. The implanted bladder had a red-burgundy color in all eleven cases. No anastomotic leakage was observed. A histological examination of the graft tissue on the 7-9 after-surgery days showed the preservation of blood flow in the tissues of the bladder and ureters. No total tissue necrosis was detected.

**Conclusions:** In our experimental model, transplantation of a donor kidney, ureters and part of the bladder to a recipient with a simulated microcyst is effectively feasible. Pigs are a relevant animal model for genitourinary organ transplantation.

**Trial Registration:** AnimalStudyRegistry.org DOI10.17590/asr.0000336. Registered 25 January 2024.

## INTRODUCTION

Dysfunction of bladder is one of the common pathologies associated with various diseases such as posterior urethral valves, neurogenic bladder, ectopic ureter or bladder exstrophy, which can ultimately lead to the development of microcystis [1, 2]

Dysuric disorders, long-term epicystostomies with microcystis contribute to the development of upper urinary tract infections and frequent attacks of pyelonephritis under the development of vesicoureteral reflux are the main causes of the development of chronic renal failure in patients with microcystis, which subsequently requires transplantation of a donor kidney. [3, 4] Donor kidney transplantation and bladder reconstruction are necessary for these pathologies.

Patients, mostly children, with neurogenic bladder often require bladder augmentation due to low volume or high intravesical pressure threatening the upper urinary tract. New reservoirs are also required for patients with bladder cancer who have undergone bladder resection. [2, 5]

Currently, in cases of microcystis with end-stage renal disease, before kidney transplantation to a potential recipient, to increase bladder capacity, improve urine storage in the bladder and urinary continence, and also to ensure protection of the upper urinary tract, augmentation or reconstructive cystoplasty operations are performed with segments of the gastrointestinal tract or other biological or synthetic tissues. [6]

After preliminary cystoplasty, the recipient with end-stage renal disease undergoes a second surgical intervention such as donor kidney transplantation. Bladder augmentation operations followed by transplantation are accompanied by technical difficulties, have a number of complications and disadvantages, such as stone formation, mucus formation, urinary infections, a lifelong need for regular catheterization with the risk of bladder perforation, intestinal obstruction, metabolic acidosis, and malignant degeneration, which often leads to early graft loss, decreased quality and life expectancy of the patients. [7]

Currently, many treatments for bladder dysfunction have been proposed, including tissue bioengineering and artificial organ, and partial bladder augmentation in animal models. [8, 9] There are a number of reports on experimental experiments on rats and mice, partial and total transplantation of a donor bladder on a vascular pedicle without a kidney and with kidneys en bloc. [7, 10]

Bladder enlargement using combined kidney and bladder transplantation with long-term survival in world practice has not yet been fully studied. [11–13]

Data on bladder transplantation in humans are very limited. Some limited studies published reports of en bloc transplantation of both kidneys and a vascularized bladder segment from pediatric donors to pediatric recipients with good functional results. [13, 14] There is also a single experience of transplanting both kidneys and part of the bladder from a pediatric donor to an adult recipient. [15] The purpose of our study was to evaluate the effectiveness of en bloc transplantation of a donor kidney, ureters and part of the bladder to a recipient with simulated microcystis in an experimental study.

## METHODS

### Participants

A total of ten White Landrace sibling pigs were used in the study.

### Placement and adaptation of the animals

The animals were delivered from a special livestock farm (Aktobe region, Kazakhstan), with which West Kazakhstan Medical University named Ospanov Marat has an agreement on raising and supplying pigs for experimental work. The animals were from the same sow and were siblings to each other in order to avoid incompatibility in the AB0 system and ensure immunological tolerance between the donor and recipient. The animals were 3-4 months old, weighing from 35 to 45 kg. The animals were housed in a specially designated room of a vivarium building at the University with an area of 14.5 m², air humidity 55-65%, room temperature from 19 to 22 □. Straw was used as bedding. Before intervention, donors and recipients were kept separate from each other. After intervention, during the observation period and before euthanasia, each animal was kept separately within auditory accessibility from other animals.

The animals were divided into 12 donors and 12 recipients. Animals were fed commercial pig feed (combined feed ASIAMIX, Hermes LLP, Republic of Kazakhstan) twice daily at an appropriate amount for growing pigs, and water was available according to their needs. The duration of acclimatization for all animals ranged from 8 to 14 days.

### Study Design

A 29-day, open, pilot prospective experimental trial: 14 days constituted an adaptation period, 5 days for the interventions, and 10 days constituted an observation period. Every day we performed one pair of operations, on the donor and the recipient. **(Figure 1)**

**Figure 1.**
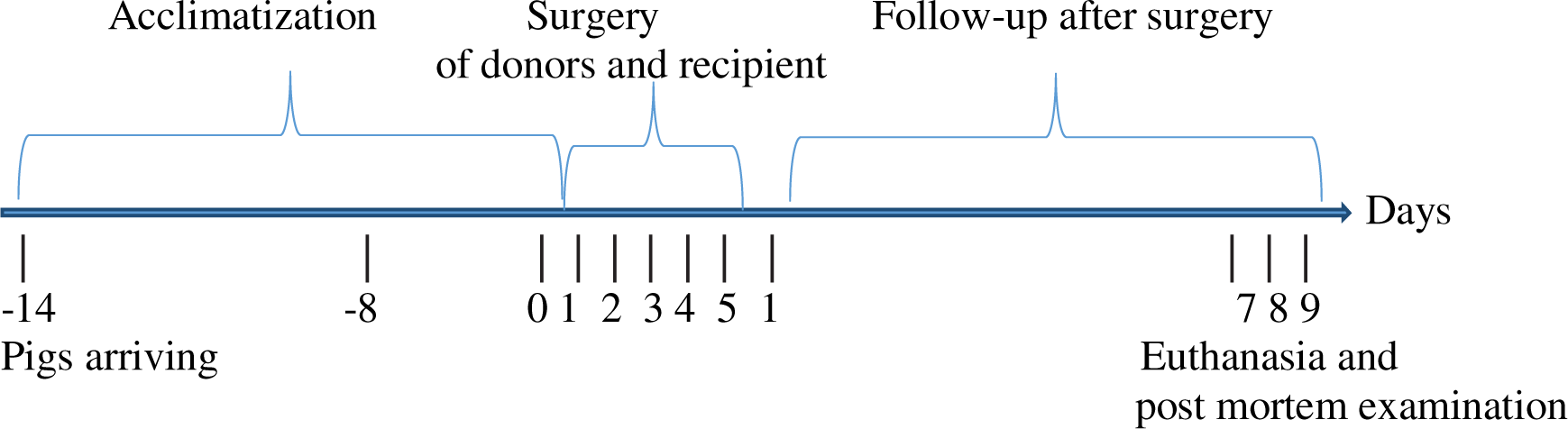
Time line of the experimental study.

### Ethic

The Local Ethics Committee of West Kazakhstan Medical University named Ospanov Marat (phone: +7-7132-543707; https:// https://zkmu.edu.kz/) approved the animal study titled “Development of simultaneous transplantation of donor kidney, ureters and part of the bladder to the recipient with simulated microcystis in the experiment on animals” (approval protocol number is #8 of 20.10.2023). The committee confirms that all methods were performed in accordance with the Animal Welfare Act (AWA) and The Public Health Service Policy on Humane Care and Use of Laboratory Animals (PHS Policy).

### Preoperative Preparations and Anesthesia

Animals were fasted for 8–12 hours and deprived of water for up to 4 hours before administration of anesthetic and analgesic drugs. Sedation was carried out directly in the animal’s enclosure. An infusion line on a needle was inserted intramuscularly into the back of the animal’s neck, and xylanite at a dose of 2 mg/kg body weight, ketamine at a dose of 7.5 mg/kg, and atropine at a dose of 0.01 mg/kg body weight of the animal were administered. After sedation, a peripheral vein was catheterized (butterfly catheter, size 22G) on the outer surface of the auricle and fixed with suturing. Loss of consciousness and relaxation occurred within 15-20 minutes. Monitoring of vital functions such as pulse oximetry, ECG cardiac monitor Life Scope TR BSM-6301K Nihon Kohden was connected, and external respiration was monitored.

Before tracheal intubation, anesthesia is deepened by administering propofol 100 mg (2.5 mg/kg) IV slowly to prevent hemodynamic disturbances. Tracheal intubation in the supine position with the head thrown back by Miller type blade #4, endotracheal tube #7.0 with a cuff, and put on a rigid guide. After intubation, a respiratory ventilator was connected, the position of the tube was controlled by comparative auscultation of the lungs and capnography on a cardiac monitor. Muscle relaxation with ardouane (pipecuronium bromide) at an initial dose of 0.05 mg/kg and a maintenance dose of 0.02 mg/kg every hour. Ventilation was performed using a Mindray SV300 device in VCV mode with ventilation parameters: tidal volume of 280 ml (7 ml/kg), RR 28/min, FiO2 50%, PEEP 5 cmH2O. The main anesthesia was total intravenous anesthesia (TIA) based on propofol and ketamine, which were administered as a continuous infusion using a Perfusor Compact C B/Braun syringe dispenser, propofol 5 mg/kg/h, ketamine 2 mg/kg/h. Nonsteroidal anti-inflammatory drug (NSAIDs) such as ketonal was used for additional pain relief.

Before the operation, catheterization of the internal jugular vein was performed with a 7F catheter, catheterization of the carotid artery with a 22 G cannula for invasive blood pressure monitoring. Venesection in the neck with fixation of catheters to the skin was used to access the great vessels.

### Intervention

#### Removal of the left kidney, ureters and bladder

On the operating table with the donor in the supine position, after fixing the limbs and shaving the surgical field, the patient was washed with water and soap, then treated with betadine solution, and covered with surgical linen. A cross-shaped incision was made in the anterior abdominal wall from the sternum to the pubis. The large and small intestines were mobilized and retracted proximally to expose the lower pole of the kidneys, the inferior vena cava, and the abdominal aorta. In an acute manner, the kidneys were completely isolated from the capsule and surrounding tissues, and the ureters and bladder were mobilized. Removal of the left kidney with the ureter, as well as part of the ureter at the level of the lower third of the right kidney and the bladder as en block was carried out after intravenous administration of heparin 10,000 units. The supra- and infrarenal aorta and inferior vena cava on each side were clamped to prevent bleeding. In the left kidney, the vein and artery were excised at the level of the inferior vena cava and aorta, respectively. The removed organs were immediately placed in a sterile bag and a cup with ice. The operation was completed in 35 minutes. The donor was euthanized by intravenous administration of 4% potassium chloride solution.

#### Back table

On a separate table, the renal artery was perfused with cold Ringer’s solution to 4-5°C until it was completely cleared of blood. 1/3 of the bladder in the transverse direction was excised and removed, leaving the area with the ureteric orifices. Then the organs were stored in a perfusion solution at a temperature of 4-5°C for no more than 40-50 minutes. (**Figure 2**)

**Figure 2.**
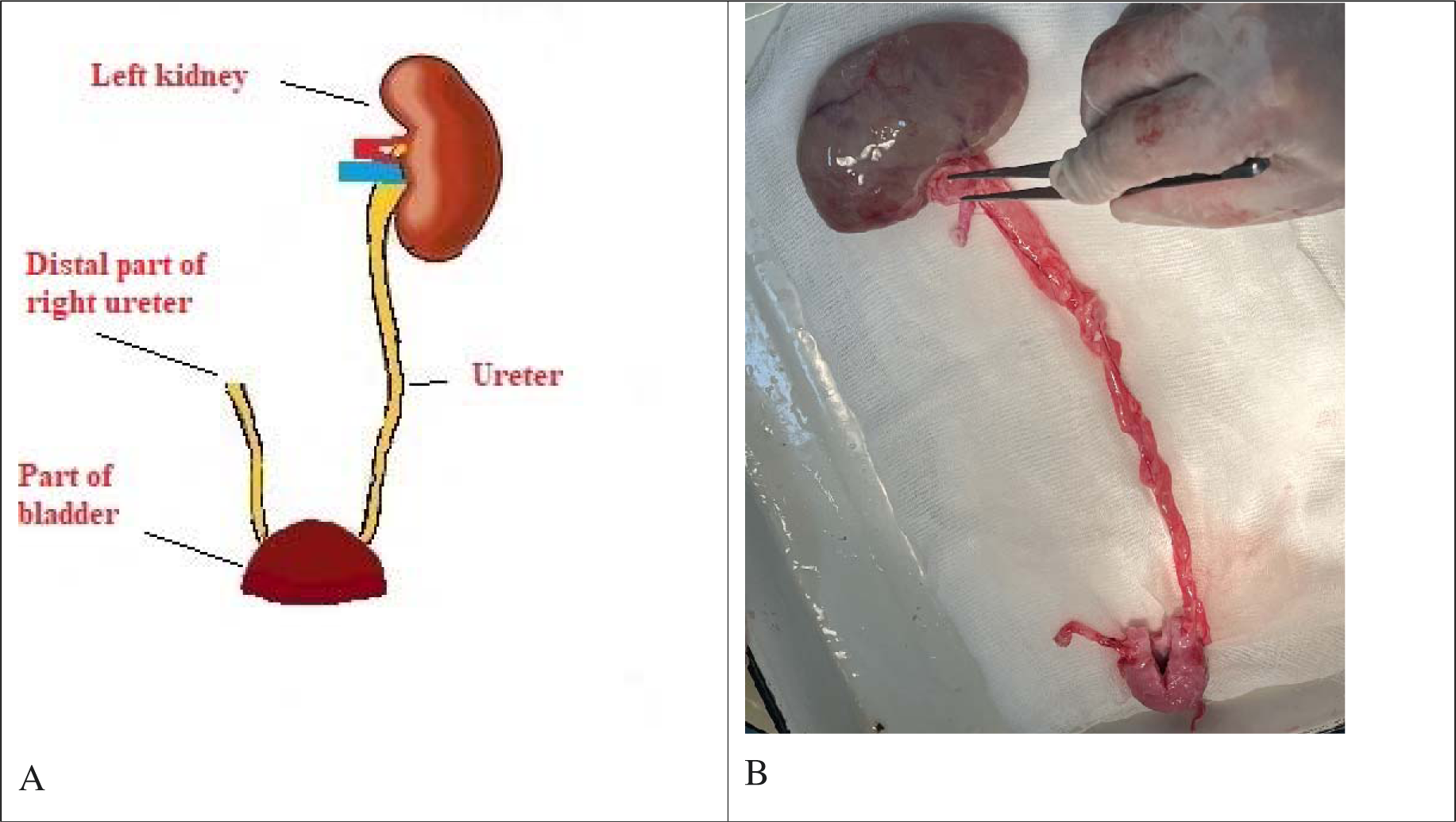
Removed from a donor organs; the left kidney, the left ureter, part of the bladder and the distal part of the right ureter: A, schematic image; B, original image.

#### Graft implantation into a recipient

Before implantation of donor organs, 30-40 minutes before reperfusion, a solution of methylprednisolone 10 mg/kg was administered intravenously as an immunosuppressant into a recipient. The recipient underwent a median laparotomy along the linea alba at a distance of 5-6 cm from the sternum to the pubis. A right-sided nephroureterectomy, ligation and dissection of the left ureter with ligation of the distal part at the level of the lower 1/3 were performed. The proximal portion of the ureter of the native left kidney was left open for subsequent anastomosis. The donor’s kidney was implanted orthotopically in place of the removed kidney. After mobilizing the inferior vena cava and clamping the vessel wall with a Satinsky clamp, an end-to-side venous anastomosis was performed with a continuous suture of 2 threads (Prolen 6/0). Then an arterial anastomosis was performed between the graft artery and the remaining end of the artery of the removed kidney in an “end-to-end” manner using a continuous suture of 2 threads (Prolen 6/0). Reperfusion and warm water dousing of the kidney were performed. After mobilization of the recipient’s bladder, half of the bladder was dissected in a transverse direction to simulate microcystis. Implantation of part of the bladder graft onto the remainder of the native bladder was performed using a continuous suture (Vicril 4/0) in 2 rows. An end-to-end ureteroureteroanastomosis was then performed between the native ureter of the left kidney and part of the graft ureter (**Figure 3**). The anastomosis was performed after installing a ureteral stent (type DJ, size 4.8Ch, length 24 cm) with a continuous suture of 2 threads (PDS 6/0). Hemostasis. Drainage of the abdominal cavity was carried out with a drainage tube 20 Ch. Interrupted sutures were placed on the wound layer by layer. Checking hemostasis. Drainage of the abdominal cavity was carried out with a drainage tube 20 Ch. Interrupted sutures were applied to the wound in layers.

**Figure 3.**
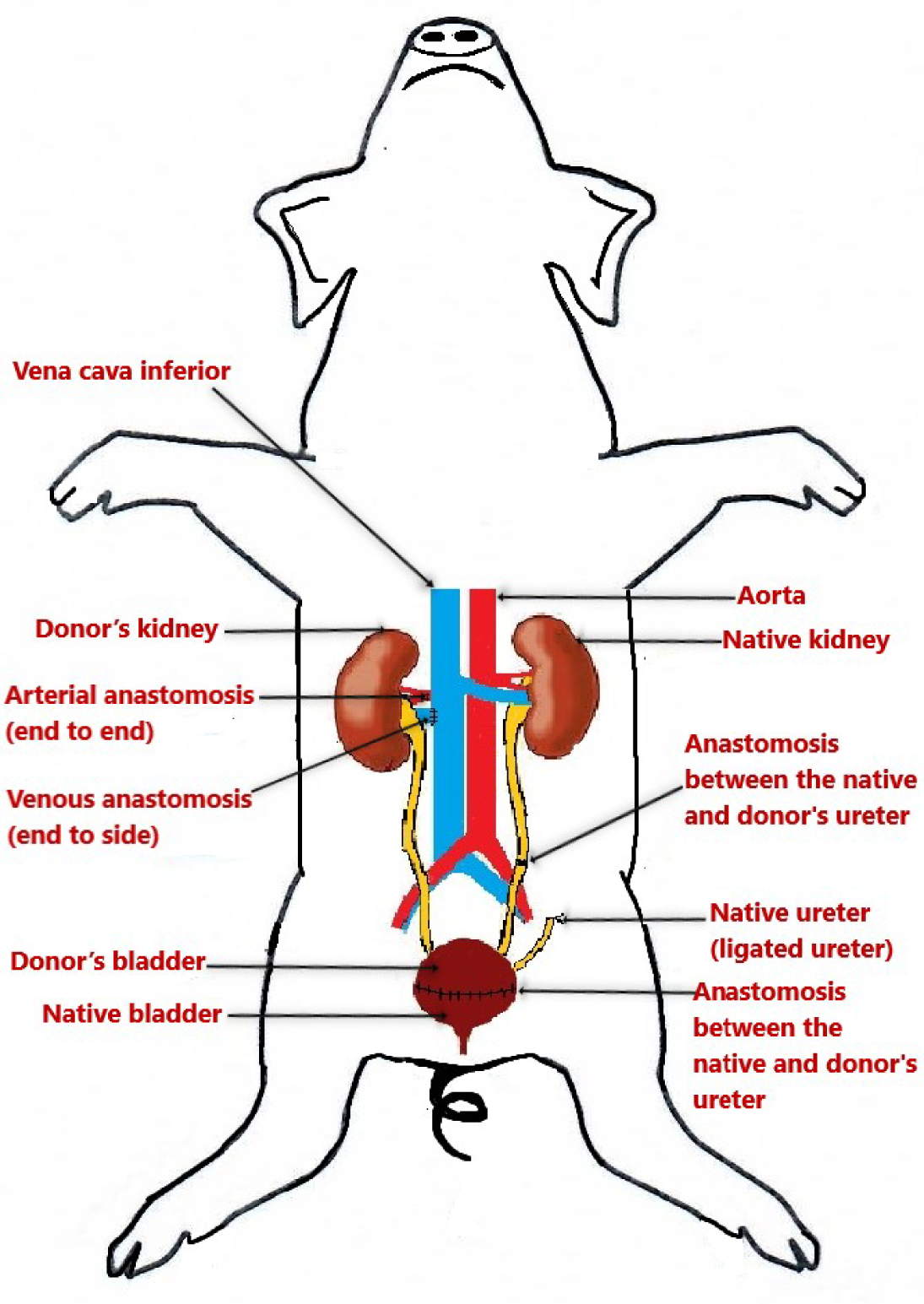
The left donor kidney, ureter, part of the bladder with the lower 1/3 of the right ureter were transplanted orthotropic as en block in place of the recipient’s removed native right kidney, right ureter, part of the bladder and the lower 1/3 of the left ureter.

After the end of the intervention, we waited for the pigs to awaken and restore muscle tone and spontaneous breathing. After turning off the respiratory ventilation, spontaneous breathing through the endotracheal tube without oxygen was assessed. In the absence of episodes of desaturation, tachy/bradypnea, and stable hemodynamics, tracheal extubation was performed.

#### Postoperative follow-up

All recipients after transplantation were transported to an enclosure, where they were observed for 7-9 days. The duration of observation was chosen to evaluate grafts on the 7th, 8th and 9th days after surgery. A prerequisite in the early postoperative period is analgesia. In the absence of narcotic analgesics, nonsteroidal anti-inflammatory drugs were used for pain management. On the first day of the postoperative period, ketonal was administered at a dose of 100 mg intravenously every 6-8 hours, followed by an increase in the interval of administration.

#### Tissue biopsy and histology

Histological examination was carried out in the Division of Pathomorphology of Department of Histology at the West Kazakhstan Medical University named Marat Ospanov (Aktobe, Kazakhstan). The obtained biopsy materials were fixed in a 10% formalin solution for 24 hours and subjected to standard histological processing and embedded in paraffin. Using an Accu-Cut SRV 200 rotary microtome (Sacura Finetek, USA), sections 3–5 µm thick were prepared from paraffin blocks and stained with hematoxylin-eosin for survey microscopy. The study of histological preparations was carried out using an Axio Lab Al microscope; photographing of the preparations was carried out with an AxioCam ERc5s digital camera.

For histological examination, tissues were taken at different periods of the operation and from different places of the graft:

1) after perfusion of the graft with Ringer’s solution during the back-table procedure:

- all layers of the bladder wall;
2) after the graft is connected to the general bloodstream:

- all layers of the bladder wall;
3) after relaparotomy, before euthanasia:

- renal graft parenchyma;
- ureter at the level of the upper 1/3, middle 1/3 and lower 1/3;
- contralateral ureter-graft
- all layers of the bladder wall;
- at the level of the anastomosis of the bladder with capture of the native and transplanted parts.

#### Relaparotomy and euthanasia

The pigs were euthanized at the end of the experiment. Before euthanasia, on days 7-8-9 after surgery, relaparotomy was performed intramuscularly under general anesthesia, visual and palpatory assessment of the transplanted organs (coloring, turgor, filling of cavities with urine), then resection biopsy of the graft tissue. Euthanasia was carried out by intravenous injection of a 4% potassium chloride solution.

### Statistics

Descriptive statistics and calculation of percentages were used. Categorical variables were presented as counts and percentages. Analyses were performed on an intent-to-treat basis.

## RESULTS

### Post mortem examination

Thus, en bloc transplantation of the kidney, ureter and part of the bladder were performed from 12 donors to 12 recipients, and the macrovisual and histological results of operations performed after 7, 8, and 9 days. **Table 1** shows the recipient survival results.

**Table 1.** Survival results of recipients (n=12) after en bloc transplantation of the kidney, ureter and part of the bladder (intention-to-treat approach)

| Animals | Survival, day/hour | Cause of death |
| --- | --- | --- |
| Recipient #1 | 10 hour | Pulmonary embolism |
| Recipient #2 | 9 days | Euthanasia |
| Recipient #3 | 8 days | Euthanasia |
| Recipient #4 | 7 days | Euthanasia |
| Recipient #5 | 7 days | Euthanasia |
| Recipient #6 | 9 days | Euthanasia |
| Recipient #7 | 8 days | Euthanasia |
| Recipient #8 | 8 days | Euthanasia |
| Recipient #9 | 9 days | Euthanasia |
| Recipient #10 | 9 days | Euthanasia |
| Recipient #11 | 9 days | Euthanasia |
| Recipient #12 | 8 days | Euthanasia |

During the present experiment, our staff spent a lot of time in the post-surgery pens observing the pigs. After the operation, the pigs were very tired and did not move much, so they lay in the same position for several hours unless the staff moved them. After anesthesia, the animals became more active and already on the 2^nd^ day moved around the pen without suffering from pain. The pigs were given food 3 times a day and water in unlimited quantities.

*Recipient #1* died 10 hour after the en-block transplantation, what happened due to a technical error in the introduction of anesthesia.

### Results of histological examination of transplanted organs at different periods of surgery

#### Recipients #2, 6, 9, 10, and 11

##### A) The kidney parenchyma on the 9^th^ day after transplantation

Minor swelling in the capsule lumen is noted in some glomeruli. The epithelium of the proximal tubules is in a state of pronounced hydropic degeneration, the lumen of the distal tubules is sharply expanded, and vascular congestion is noted (**Supplemental figure 1**).

##### B) The ureter of the kidney transplant on the 9th day after transplantation

The structure of the ureter is preserved. The vessels are full of blood in all layers. Swelling and hemorrhage are noted in the outer shell. Some cells of the transitional epithelium of the mucous membrane are vacuolated, and moderate swelling and hemorrhage are noted in the muscular layer (**Supplemental figure 2**).

The structure of the ureter is preserved. Swelling and hemorrhage of the wall are noted. The vessels are full of blood in all layers. Vacuolization of the transitional epithelial cells of the mucous membrane is noted, and moderate swelling and hemorrhage are noted in the muscle layer. (**Supplemental figure 3**).

##### D) The transplanted bladder on the 9th day after transplantation

On a specimen of the bladder, the transitional epithelial cells of the mucous membrane are vacuolated, the vessels are full of blood (**Supplemental figure 4**).

#### Recipients #3, 7, 8, and 12

##### A) The kidney parenchyma on the 8th day after transplantation

Some glomeruli are reduced in size. The epithelium of the proximal tubules is in a state of hydropic degeneration, the lumen of the distal tubules is moderately expanded in certain fields of view. Foci of lymphocyte-like infiltration are noted in the stroma. Stasis of capillaries, the lumen of some vessels is empty, some are full of blood (**Supplemental figure 5**).

##### B) The ureter of the kidney transplant on the 8th day after transplantation

The general structure of the ureter is preserved, but the transitional epithelium of the mucous membrane is desquamated in places, the vessels are full-blooded in all layers, hemorrhage and moderate edema are noted in the muscular and outer adventitia (**Supplemental figure 6**).

##### C) The transplanted bladder on the 8th day after transplantation

The epithelium is thinned, desquamated, the muscle fibers are moderately loosened, there is pronounced swelling, hemorrhage and moderate inflammatory infiltration, the vessels are dilated and full of blood (**Supplemental figure 7**).

##### D) The contralateral graft ureter on the 8th day after transplantation

The structure of the ureter is preserved. Swelling and hemorrhage of the wall are noted. In all layers the vessels are full of blood. Vacuolization of the transitional epithelial cells of the mucous membrane is noted; moderate swelling and hemorrhage are noted in the muscle layer (**Supplemental figure 8**).

#### Recipients #4 and 5

##### A) The kidney parenchyma on the 7th day after transplantation

In some glomeruli, slight swelling is noted in the lumen of the capsule. The epithelium of the proximal tubules is in a state of pronounced hydropic degeneration, the lumen of the distal tubules is sharply expanded, there is congestion of the vessels.

##### B) The ureter of the kidney transplant on the 7th day after transplantation

The structure of the ureter is preserved. The vessels are full of blood in all layers. Swelling and hemorrhage are noted in the outer shell. Some cells of the transitional epithelium of the mucous membrane are vacuolated, and moderate swelling and hemorrhage are noted in the muscular layer.

##### C) The contralateral graft ureter on the 7th day after transplantation

The structure of the ureter is preserved. Swelling and hemorrhage of the wall are noted. The vessels are full of blood in all layers. Moderate swelling and hemorrhage are noted in the muscle layer.

##### D) The transplanted bladder on the 7th day after transplantation

On a specimen of the bladder, the transitional epithelial cells of the mucous membrane are vacuolated, the vessels are full of blood.

## DISCUSSION

The present study confirmed that pigs can be a pertinent animal model for genitourinary organ transplantation research. The advantage of using pigs as models is that the pig urological system is more similar to the human than most other animal models, both in aspects of physiology and anatomy. [16, 17] Pigs are also relatively cheap and reproduce quickly.

In the present study, 12 pigs weighing from 35 to 45 kg were used, regardless of gender. The experiment included offspring from the same sow to avoid transplant rejection.

Acclimation period of the animals was successful, demonstrating the importance of including such a period in most pig procedures to prevent stress. The authors recommend that pigs be delivered two weeks before the start of the experiment for successful acclimatization so that the pigs become accustomed to the food and environment. [18, 19]

In this experiment, the animals were divided into 2 groups, 12 donors and 12 recipients. The donors’ left kidney, ureter and part of the bladder were removed along with the distal end of the right ureter as en block. Then the removed organs were implanted into the recipient with preliminary removal of the recipient’s right kidney, ureter and part of the bladder with the distal part of the right ureter.

The anesthesia protocol allowed for better pain relief before and during surgery. We decided not to use muscle relaxants, as they can mask the pain during the procedure. During the operation, there were some critical points when the pigs’ blood pressure dropped, which included: reperfusion of the transplanted kidney, nephrectomy of the native kidney, and keeping the intestines out of the body. From all pigs the first pig died 10 hours after the operation that happened due to a technical error in the introduction of anesthesia. According to the pathological autopsy, the cause of death was pulmonary embolism. As other experiments have shown, intraoperative complications are not uncommon. [20, 21]

Visually, in all euthanized cases, the kidney, ureters and bladder transplanted as en block were filled with urine, full of blood, and tissue turgor was good. Coloring during visual inspection of the kidney and ureters was satisfactory, bright red. The implanted bladder had a red-burgundy color in all 11 cases. No anastomotic leakage was observed. A histological examination of the graft tissue on the 9th, 8th and 7th days after surgery showed the preservation of blood flow in the tissues of the bladder and ureters. Overall, no total tissue necrosis was detected.

In the present study, we kept pigs under anesthesia for a longer period of time and showed that organ transplantation can be performed in a manner similar to the procedure performed in humans.

It can be said that en block transplantation of a kidney, ureters and part of the bladder in an experiment on pigs with simulated microcystis is quite possible and, with qualified surgical techniques, it can be transferred into clinical transplantation with a good probability of success in humans with terminal kidney lesions in combination with microcystis.

Current clinical strategies for bladder reconstruction or substitution are associated to serious problems. [22] Our results showed that engraftment of part of the bladder as a graft en bloc with the ureters and kidney is quite possible with adequate techniques for removal, perfusion and implantation.

Postoperative care is of paramount importance. In the postoperative period, it is recommended to keep pigs separately from each other, since surgical wounds can be attacked by other pigs. Therefore, the pigs in the present study were kept in separate rooms within auditory proximity of each other to stimulate their social needs. Anesthesia after the intervention is recommended for the animals to become more active.

Adams et al. [23] showed that it is possible to use tacrolimus and mycophenolic acid as immunosuppressive therapy in pigs. Since all individuals were from the same sow, in the present study we used methylprednisolone as induction immunosuppressive therapy; graft rejection was minimal. If the pigs were from different parents, then rejection would be inevitable. [24] Rejection can be seen histologically on the third day after surgery and chemically on the fourth day. [16, 25, 26] In the present study, the pigs lived 7-9 days after transplantation before they were euthanized. The pigs were euthanized before death to allow visual and histological evaluation of the en block transplanted organs. The present experiment showed promising results, and we now plan to repeat this study in perspective to confirm the results to more reliably determine the survival of such a graft.

### Limitations

The strengths of this study are that we evaluated the effectiveness of en bloc transplantation of a donor kidney, ureters and part of the bladder to a recipient with simulated microcystis in an experimental trial. The limitations of this study are that it was a single-center experimental study, it was short in duration, it did not include T-cell/B-cell subpopulation data, and it was a small number of animals for study. Further studies with involving a large number of animals, long-term follow-ups are needed to confirm and extend the results of the study.

## CONCLUSIONS

Thus, en bloc transplantation of a donor kidney, ureters and part of the bladder to a recipient with simulated microcystis is effectively feasible in our experimental model. Pigs are an excellent animal model for en block genitourinary organ transplantation. After additional study it can be transferred into clinical practice for humans with end-stage kidney disease in combination with microcystis.

## DECLARATION

The study was carried out in the Republic of Kazakhstan in August-November, 2023. The study was carried out in the experimental room of a vivarium building at the West Kazakhstan Medical University named Ospanov Marat (Aktobe city) by researchers from University medical center (Astana city).

### Consent for publication

All the authors of the manuscript affirm that they had access to the study data and reviewed and approved the final version.

### Availability of data and materials

The datasets used and/or analyzed during the current study are available from the corresponding author on reasonable request and with permission of the Ministry of Education and Science of the Republic of Kazakhstan. Those wishing to request the study data should contact Principal Investigator of a research grant: Dr Gani Kuttymuratov (, phone + 7-701-8812206).

### Author contributions

*GK:* study design, data collection, interventions for animals, bibliography review, statistical analysis, data interpretation, writing of the paper. *AA* and *AM:* study design, data collection, interventions for animals, data interpretation, and paper review. *KO:* study design, data collection, data interpretation, bibliography, and paper writing. *DZh* and *ZhI:* data collection, interventions for animals, data interpretation, methods and discussion writing, and paper review. *AT* and *BA:* data collection, interventions for animals, data interpretation, and paper review. *ArA:* research management, data collection, preparation of the statistical data in Excel, and paper review. *TA:* interventions for animals, data collection, and paper review. *NU:* study design, bibliography search, research management and review. *AK:* interventions for animals, data collection and paper review. All the authors read and approved the final manuscript.

## ACKNOWLEDGMENTS

This research has been funded by the Science Committee of the Ministry of Science and Higher Education of the Republic of Kazakhstan (Grant #АР19677564).

This study was conducted jointly with employees of the West Kazakhstan State Medical University named Marat Ospanov, Aktobe. We would like to thank everyone who took part in this study.

## SUPPLEMENTAL FIGURES

**Supplemental figure 1.**
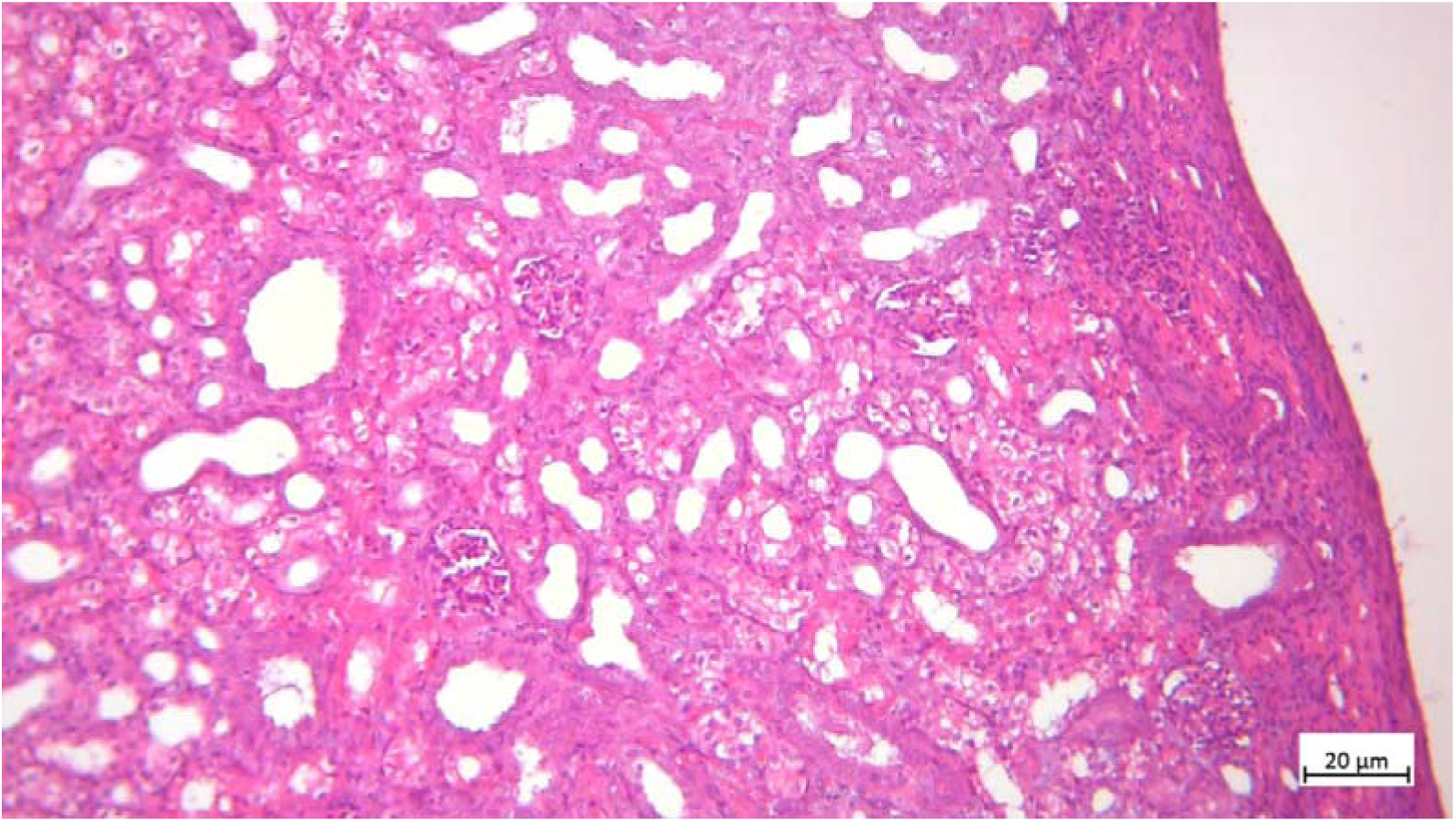
The kidney parenchyma on the 9th day after transplantation

**Supplemental figure 2.**
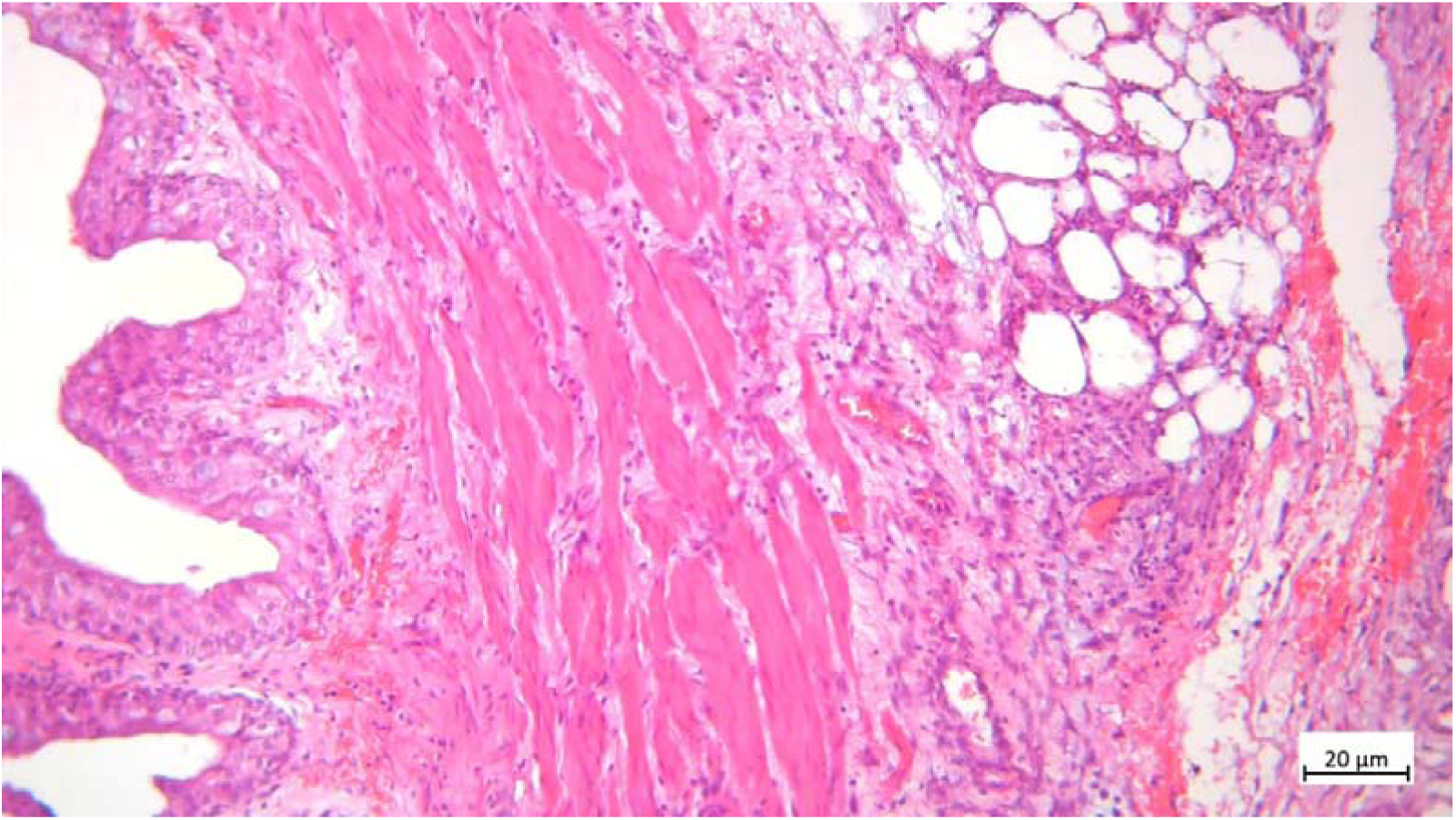
The ureter of the kidney transplant on the 9th day after transplantation C) The contralateral graft ureter on the 9th day after transplantation

**Supplemental figure 3.**
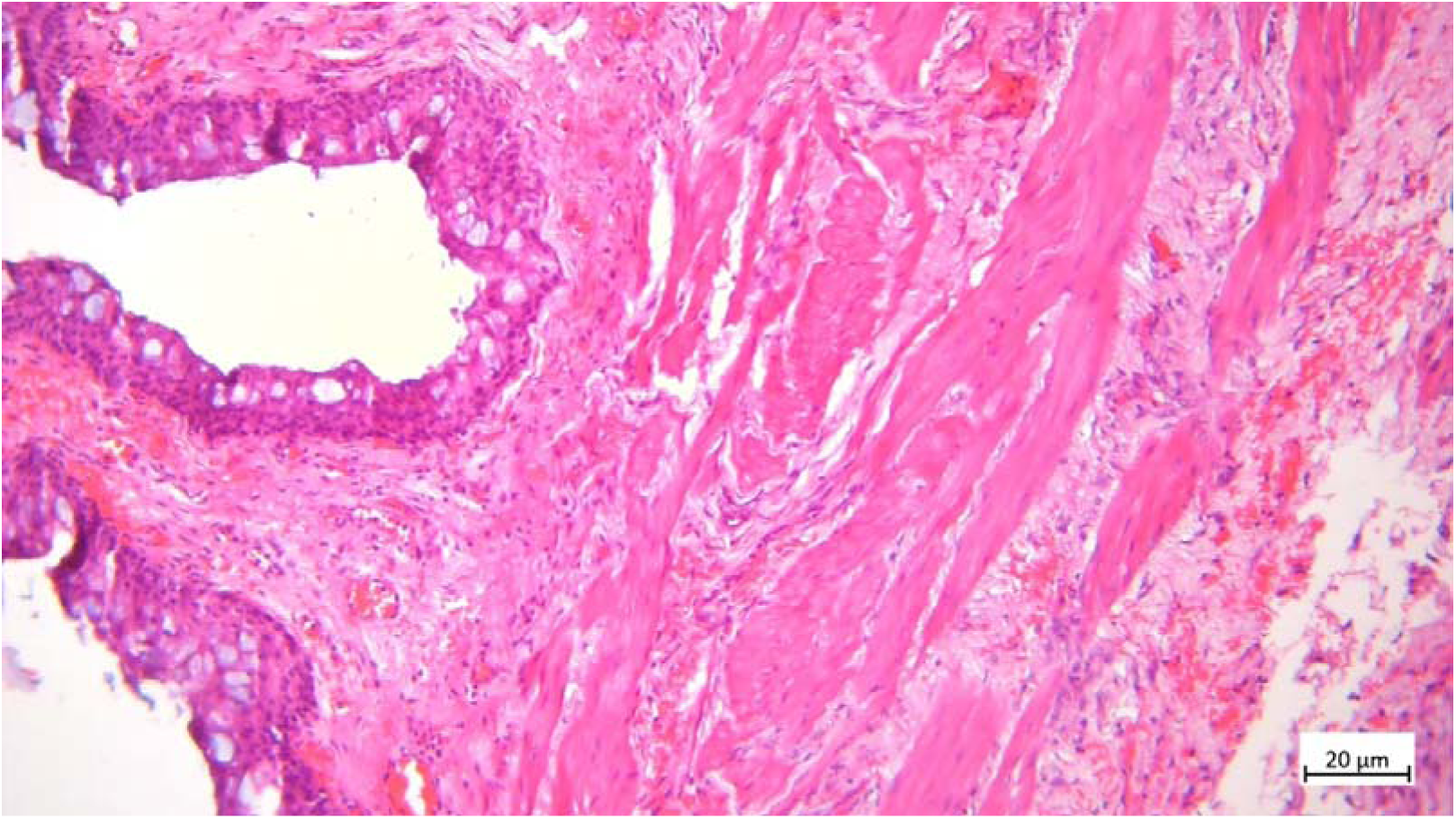
The contralateral graft ureter on the 9th day after transplantation

**Supplemental figure 4.**
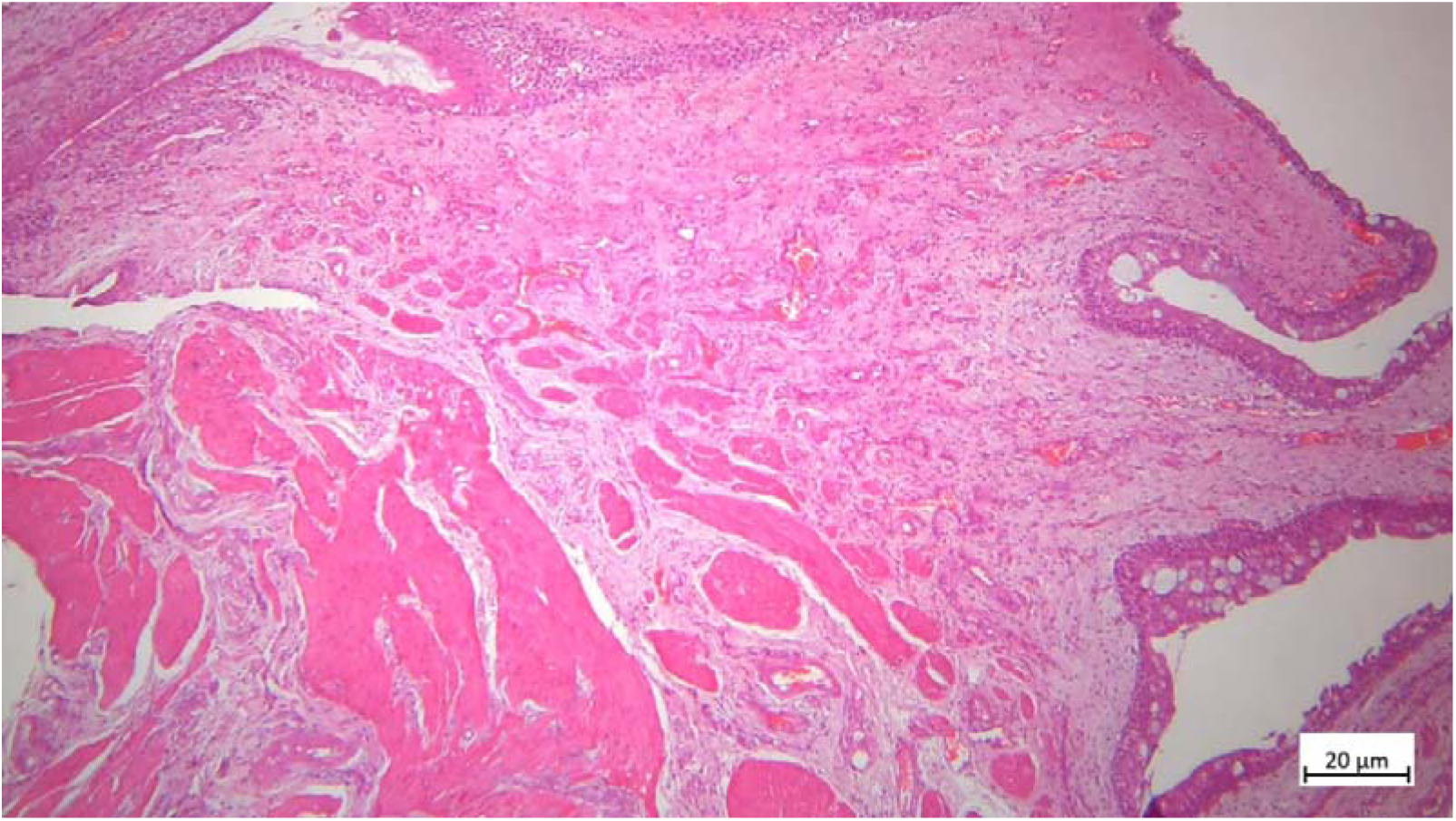
The transplanted bladder on the 9th day after transplantation.

**Supplemental figure 5.**
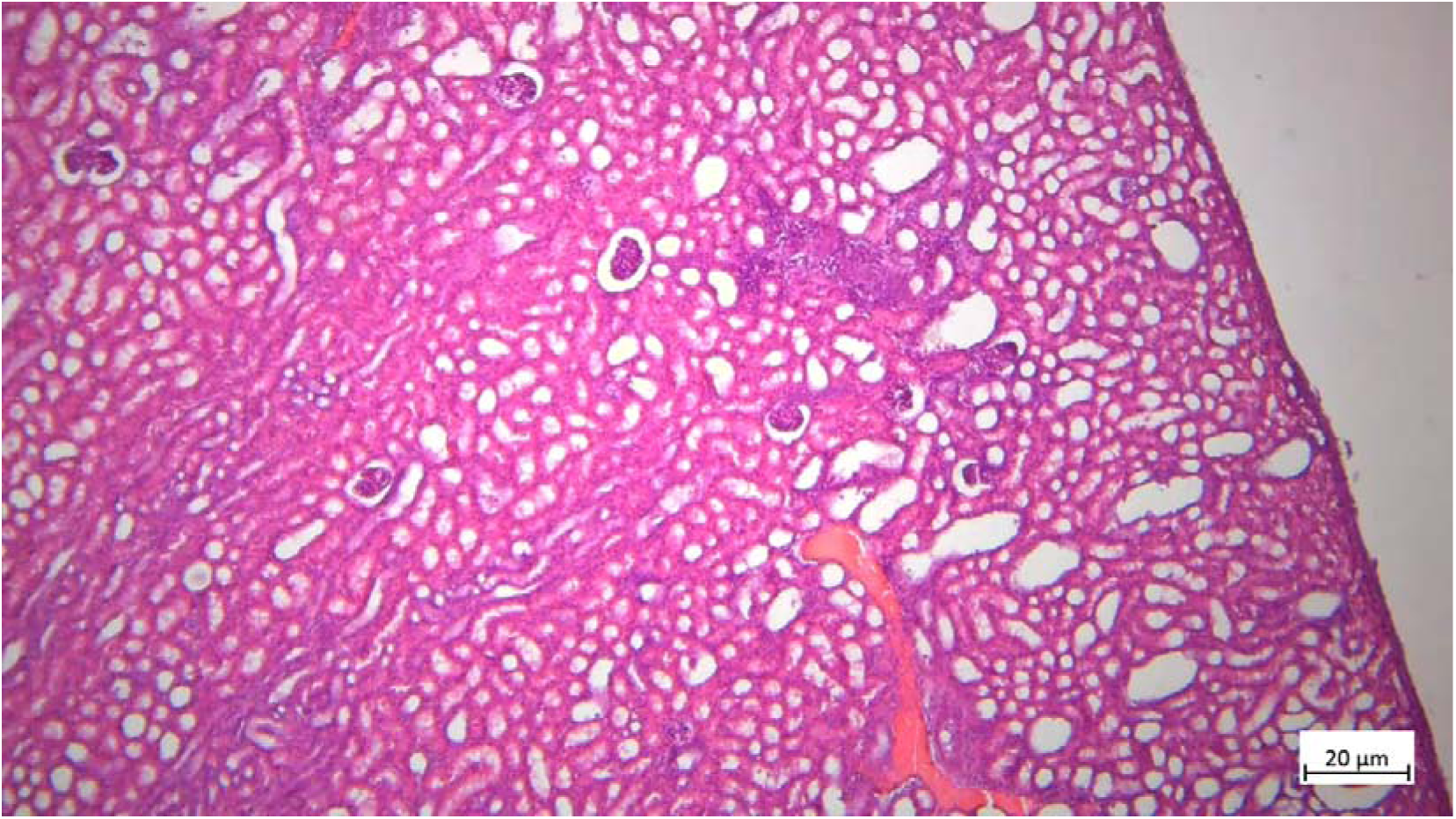
The kidney parenchyma on the 8th day after transplantation.

**Supplemental figure 6.**
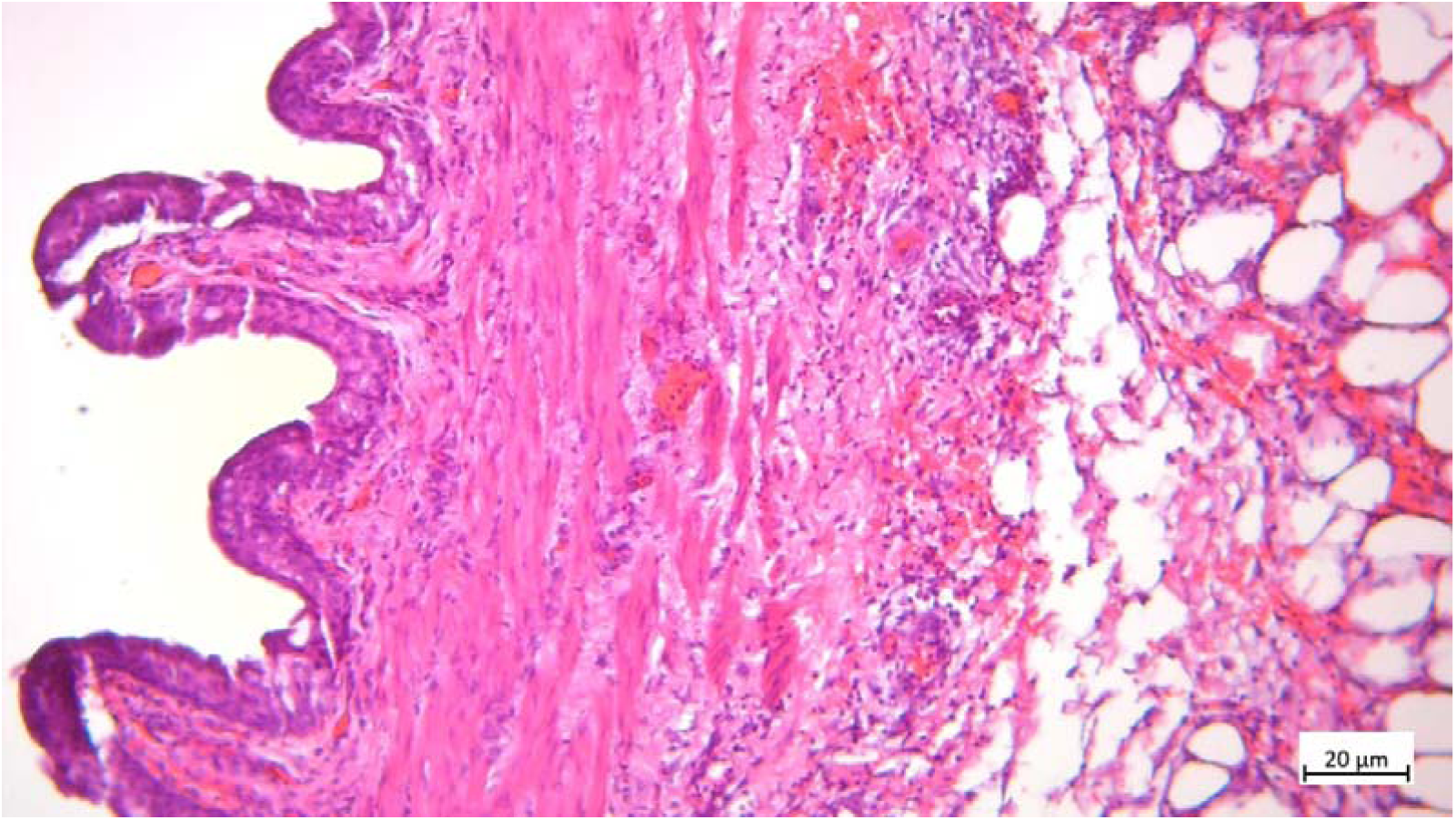
The ureter of the kidney transplant on the 8th day after transplantation

**Supplemental figure 7.**
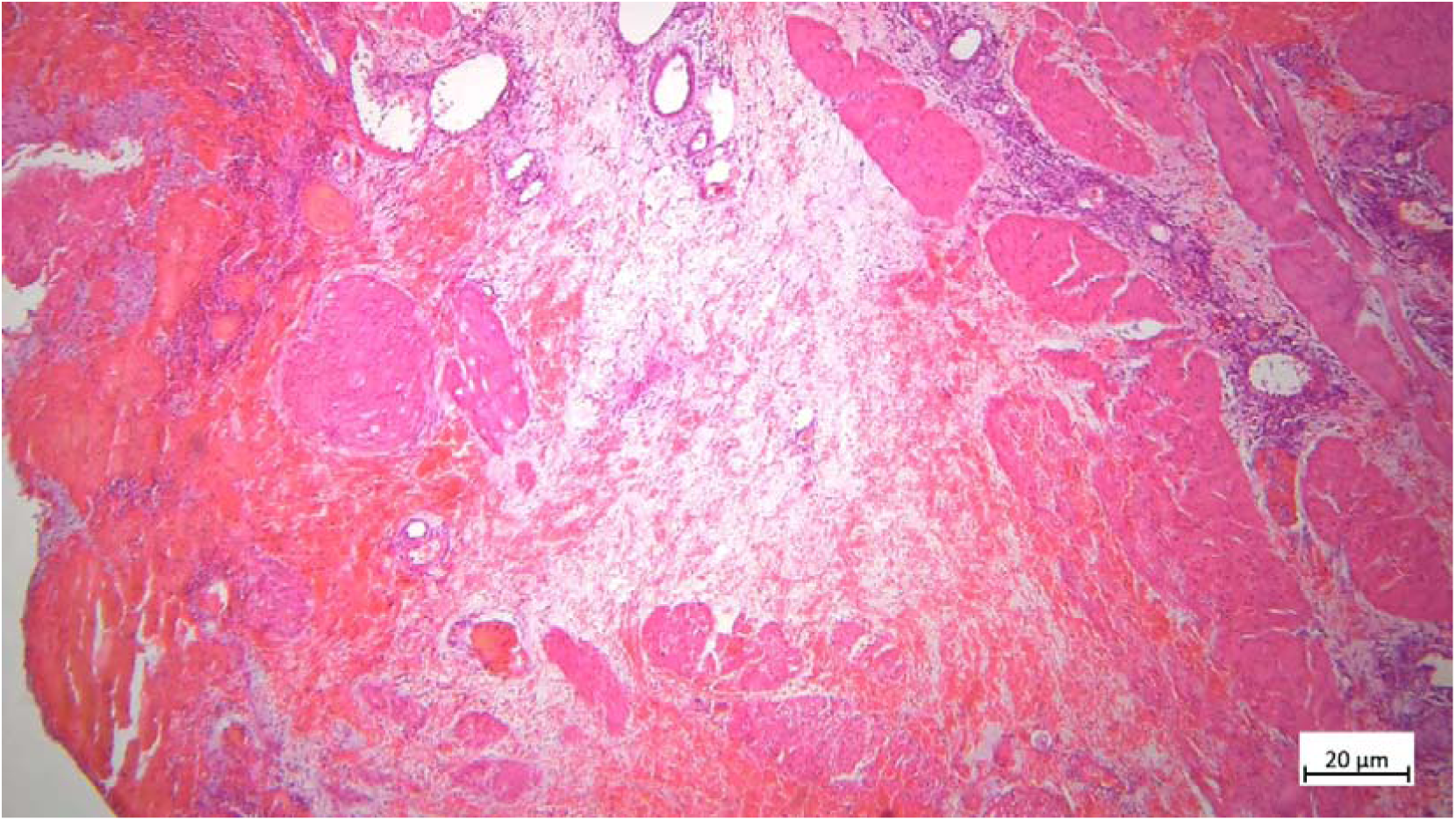
The transplanted bladder on the 8th day after transplantation.

**Supplemental figure 8.**
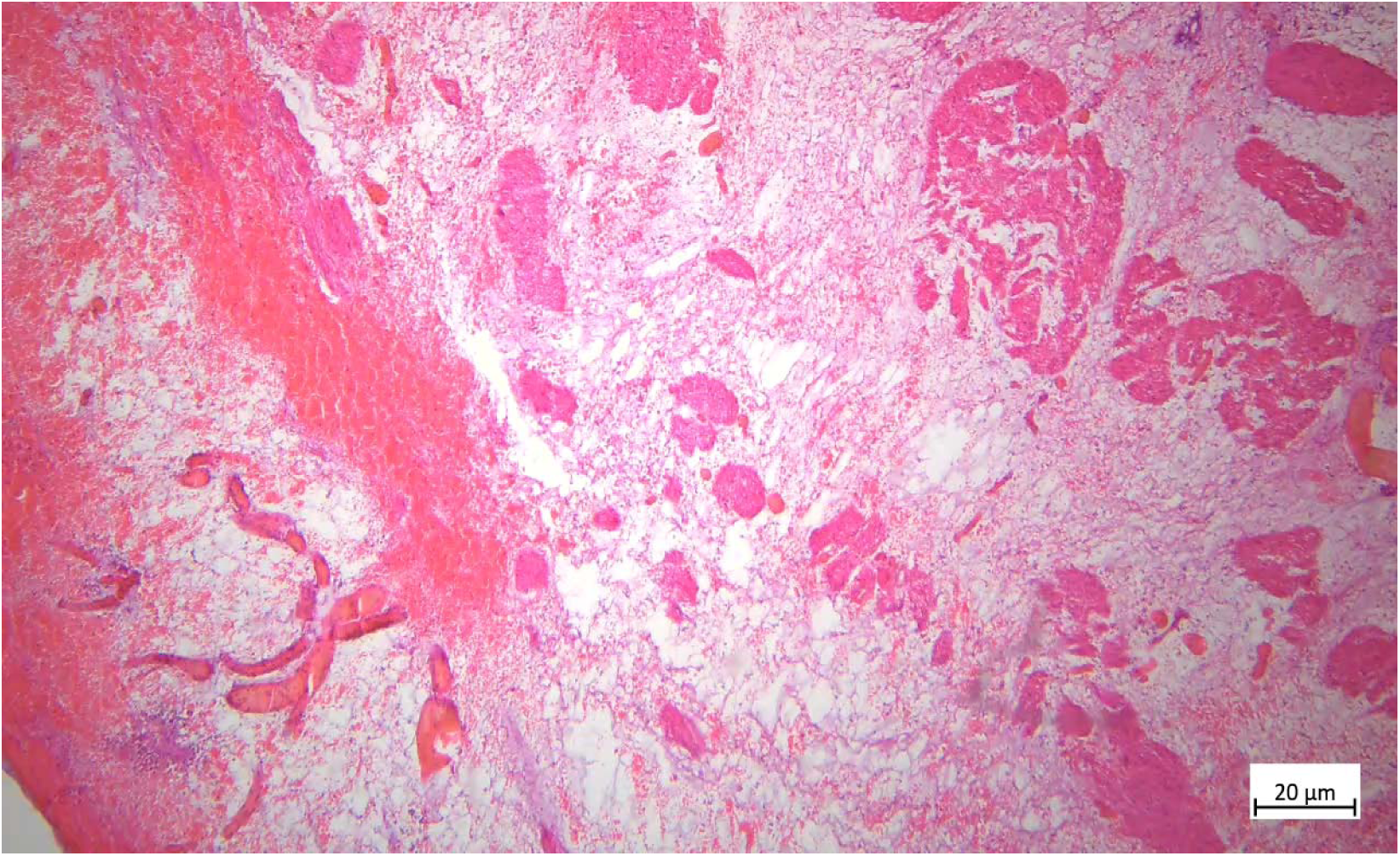
The contralateral graft ureter on the 8th day after transplantation.

